# High resolution longitudinal molecular and morphological tracking of planktonic threats to salmon aquaculture: new candidates and old players

**DOI:** 10.1101/2023.09.04.556215

**Authors:** María Algueró-Muñiz, Sofie Spatharis, Toni Dwyer, Michele de Noia, Bachar Chaib, Brendan Robertson, Calum Johnstone, Jennifer Welsh, Annabell Macphee, Marta Mazurkiewicz, Ralph Bickerdike, Hervé Migaud, Clara McGhee, Kim Præbel, Martin Llewellyn

## Abstract

Marine-phase salmonid aquaculture is a major component of the coastal economies of Northern Europe, North America and Chile and is under threat from numerous challenges to gill health, many of which originate from the phyto- and zooplankton. Associated losses are growing as a proportion of production year on year. A first step towards mitigating losses is to characterize the biological drivers of poor gill health. Numerous planktonic species have been implicated, including toxic and siliceous microalgae, hydrozoans and scyphozoans; however, rigorous longitudinal surveys of planktonic diversity and gill health have been lacking. In the current study, we present and assess an ‘exhaustive’ identification approach combining both morphological and molecular methods (environmental DNA metabarcoding) approaches in combination with robust statistical models to identify the planktonic drivers of complex gill disease (CGD) and fish mortality. We undertook longitudinal molecular and microscopic evaluation at two marine aquaculture facilities on the west coast of Scotland using daily data collected during the 2021 growing season (March-October). Examining these two different sites, one sheltered and one exposed to the open sea, we identified new, important, and unexpected planktonic drivers (e.g. doliolids and appendicularians) of CGD and mortality and confirmed the significance of some established threats (e.g. hydrozoans and diatoms). We also explored delayed or ‘lagged’ effects of planktonic abundances on gill health and undertook a comparison of environmental DNA metabarcoding and microscopy in their ability to identify and quantify planktonic species. Our data highlight the diversity of planktonic threats to salmonid aquaculture as well as the importance of using both molecular and morphological approaches to detect those. Despite our study relying on two farm sites only, our results evidence the role of the different planktonic players on salmon gill disease; there is now an urgent need to expand systematic longitudinal molecular and morphological approach across multiple sites and over multiple years. The resultant catalogue of main biological drivers will enable early warning systems, new treatments and, ultimately, a sustainable platform for future salmonid aquaculture in the marine environment.

## INTRODUCTION

Aquaculture is the fastest-growing food production sector globally and has undergone rapid expansion and diversification in recent decades (Naylor et al., 2021). Current marine production of finfish accounts for over 8.3 million tonnes globally (FAO, 2022) and is dominated by four species: Atlantic salmon (*Salmo salar*), rainbow trout (*Oncorhynchus mykiss*), European seabass (*Dicentrarchus labrax*), and gilt-head seabream (*Sparus aurata*) (Naylor et al., 2021). Most marine finfish are grown in floating cages anchored to the seabed in the coastal environment. Unlike their wild progenitors, fish in net pens are contained within the net volume and whilst they do show avoidance behavioural response to water quality changes, they have limited ability during significant phytoplankton bloom or jellyfish swarm events. Consequently, marine finfish aquaculture is heavily dependent on the quality of the local biotic and abiotic environment to sustain survival and optimal growth. Climate change is affecting aquaculture worldwide at local and global scales (Maulu et al., 2021). Coastal environments are experiencing the simultaneous hit of multiple anthropogenic stressors (Martinez & Rusch, 2021) such as warming and eutrophication; sea surface warming at an unprecedented rate, meanwhile terrestrial nutrient run-off from domestic and agricultural activities is driving coastal phytoplankton blooms (Dai et al., 2023). The resultant combination of increasing phytoplankton and zooplankton productivity, alongside accelerated parasite development and lowered dissolved oxygen (DO) (Dalvin et al., 2020; Guerrero et al., 2018; Jones & Price, 2022), represents a significant and growing challenge to aquaculture in coastal environments globally.

In recent years, poor gill health has emerged as a major contributor to reduced farmed finfish survival at sea, especially in Atlantic salmon (Boerlage et al., 2020; Herrero et al., 2022). Gill health-related losses in the Scottish salmon aquaculture have been steadily increasing particularly in the late summer months, e.g. 1.57% in September 2018 compared to 4.65% in September 2022 (Salmon Scotland, 2022). The gill plays several vital roles in teleost physiology: a mucosal barrier for the immune system, gas exchange, osmoregulation, excretion of nitrogenous waste and hormone production (Foyle et al., 2020). The gill surface is in continuous contact with the marine environment and as such continually exposed to potential biotic and abiotic stressors. Gill health challenges are multifactorial as several transmissible parasites, viruses, and bacteria are implicated (e.g. Rodger H, 2007). Nematocysts, stinging organelles associated with cnidarian epithelial cells, are also thought to drive gill damage and inflammation (Kintner & Brierley, 2019). Phytoplankton species may also have a role, either via toxin production or, in the case of siliceous diatoms, as direct damage to the gill via penetration of the siliceous spines in gill tissue (Bell, 1943; Østevik et al., 2022). Biological stressors on salmon gill health may synergistically interact to drive cumulative damage to gill tissue. For example, initial exposure to hydrozoans can result in secondary parasitic infection (e.g. Kintner & Brierley, 2019), although experimental challenge trials have been inconclusive in proving the link between hydrozoan exposure and subsequent *Neoparamoeba perurans* infection severity (Bloecher et al., 2018).

Pathological responses in salmon gills encompass amoebic gill disease (AGD), proliferative gill disease (PGD) and other gill damage that fall under the umbrella ‘syndrome’ of complex gill disorder (CGD) (Noguera & Marcos Lopez, 2019). CGD includes a wide range of clinical gill disease presentations generally occurring from summer to late autumn on marine Atlantic salmon farms and relates to both AGD and PGD, which can cause gill necrosis, respiratory distress and, ultimately, fish death. For most PGD cases in farmed Atlantic salmon, no specific pathogen or other harmful agent can be incriminated as the causative agent. Over 40 zoo- and phytoplankton species are suspected to be involved (Boerlage et al., 2020). Many salmon producers monitor the water column for these species daily to detect blooms in a timely fashion and instigate the limited mitigation measures at their disposal (e.g. feed reduction, plankton skirts and bubble nets).

Currently, the primary mode of plankton monitoring on fish farms involves morphological identification via light transmission (phytoplankton) and binocular (zooplankton) microscopy in mostly single time-point water samples taken throughout the water column. Accurate taxonomic identification of morphologically similar species is challenging and time-consuming, is subject to human error, with significant taxonomic expertise required in-house (Deagle et al., 2018). Effective disease mitigation relies on early intervention following detection above arbitrary thresholds, especially for algal and cnidarian blooms (Engehagen et al., 2021). To achieve this, planktonic monitoring must be rapid, accurate, quantitative and encompass multiple taxonomic levels including metazoans and protists. Furthermore, many of the planktonic drivers of gill pathology in salmonid aquaculture are unknown, and/or their association with poor gill health is circumstantial and lacks a robust statistical framework.

In the current study, we assessed an ‘exhaustive’ approach to plankton identification combining both morphological and molecular (environmental DNA a.k.a. eDNA metabarcoding), in combination with robust statistical models, to identify the planktonic drivers of CGD and mortality in salmonid aquaculture. To achieve this, we undertook longitudinal molecular and microscopic evaluation on a daily basis during the production period (March-October 2021) at two marine aquaculture facilities on the west coast of Scotland. Examining these two sites, one sheltered and one exposed to the open sea, we identify new and unexpected planktonic drivers of CGD and mortality and confirm the importance of some established threats. We also explore delayed or ‘lagged’ effects of planktonic abundances on gill health as well as undertaking a comparison of eDNA metabarcoding and microscopy in their ability to identify planktonic species and estimate their abundance.

## METHODS

### Sites and sample size

Between March and September 2021, we monitored two sites on the NW coast of Scotland, UK: a sheltered site with a large freshwater input was monitored for 223 sampling days, whilst an exposed open water site provided us with 191 sampling days. This daily monitoring included phytoplankton and zooplankton surveys, as well as eDNA sampling. Moreover, matching our sampling period, our collaborators in the farms provided mortality (z-scores), PGD and AGD scores, environmental data related to temperature, oxygen levels, salinity, and visibility in the water adjacent to the cages, and treatments applied to fish during our study. We then used best-fitting models to determine the strongest planktonic predictors of diminished fish health.

### Plankton sampling and identification via microscopy

Species were identified by both classic (morphological) and molecular approaches. Whilst the classic approach was thought to offer a more accurate representation of species abundance, we considered that the molecular approach might offer a more exhaustive species list—at the level of the operational taxonomic unit (OTU)—as it can capture species that are rare and/or small and thus undetected by microscopy. Samples were analysed morphologically using conventional microscopy techniques, and genetically via metabarcoding of a fragment of the cytochrome oxidase I (COI) gene (see next section).

Phytoplankton was sampled at 5m depth with a Van Dorn bottle and 250 mL of seawater were immediately preserved with acidic Lugol’s iodine solution (5 %). Samples were preserved in dark glass bottles in a dark cool place for 3 weeks to 6 weeks before microscopic analysis. Samples were then filtered through nitrate cellulose membranes of 0.45 μm mesh (Fournier, 1978) followed by transillumination of the filter with immersion oil and observation under a Zeiss Axiolab upright microscope. This approach was adopted due to the observed underestimation of the smaller fraction of phytoplankton (<20 μm) by the inverted microscope method (Utermöhl, 1958). Taxa were identified in their majority to species level and cell counts were expressed as number of cells per Liter.

Zooplankton was sampled in the vicinity of the salmon nets by vertical net hauls with an Apstein net (55 μm mesh size, 25 cm diameter) equipped with a closed cod end. Sampling depth was restricted to 10 m to mimic the depth of the fish net pens. During the experimental period, one net haul per site was collected, every net haul consisting of a total filtered volume of 491 L. Samples were then rinsed with seawater prefiltered through 50 μm mesh, collected in containers and preserved in 4 % formaldehyde buffered with sodium tetraborate. During analysis, organisms were sorted using a stereomicroscope (Leica S9i) and classified to the lowest possible taxonomic level.

Initial target zooplankton taxa were represented by salmon sea lice and hydromedusae, based on literature references and direct communication with the farms. Copepod nauplii from different species were pooled together, except for sea lice species *Lepeophtheirus salmonis*, which were monitored independently. Every sample was sieved through 50 μm mesh, rinsed with tap water and poured into a calibrated beaker, where organisms were well mixed before subsampling three aliquots with a Hensen Stempel pipette (Harris, 2000), representing a minimum of 12 % volume of the sample. Counting was restricted to the 12 % of volume for the most abundant taxa, whereas the remaining sample volume was monitored for the taxa not recorded in the aliquots to record diversity, with special focus on the target species.

### Environment DNA sampling, metabarcoding and sequencing

Water from 5 m depth was sampled by using a Niskin bottle (Sheltered site) or a Van Dorne (Exposed site), and 500 mL of seawater was filtered through a sterile 0.2 μM filter (Sterivex, Merck) in technical duplicate on-site by aquaculture staff using a modified Spear & Jackson 5 L pump sprayer. The sprayer was rinsed thoroughly in seawater each day and 1 L of sampled water used to flush the system prior to connecting the filter unit. Filters were pumped dry, sealed into sterile 50 mL centrifuge tubes including c.20 g of silica bead desiccant and stored at room temperature prior to transfer to the molecular biology laboratory in weekly batches via courier. DNA extraction was achieved using a Qiagen DNEasy Blood and Tissue® kit following the manufacturer’s protocols with a minor modification. As a first step, 500 μL of lysis buffer was added to each filter unit, and the unit was gently shaken overnight at 56℃ on a rotary wheel prior to transferring the lysate to the spin-column, washing, and elution into a 2 mL centrifuge tube as in (Turon et al., 2022). Negative extraction controls were included in the lab following the same protocol. The partial COI Leray-XT fragment (313 bp) was amplified from these metagenomic DNA samples using the mlCOIintF-XT/jgHCO2198 primer pair (Wangensteen et al., 2018). Leray primers included spacers to increase complexity, as well as dual index barcodes to assist with the multiplex. PCR products were bead-purified prior to PCR-based tagging to achieve high-level (600+) multiplex using dual index 96 barcoded primers, alongside Illumina P5 and P7 adaptor and Nextera sequencing primer binding sites (see supplementary data for primer sequences). The final library was gel purified, quantified and submitted to 250bp paired-end sequencing on an Illumina NovaSeq 6000 instrument at Novogene PLC.

Custom demultiplexing of internal and external barcodes was undertaken using a combination of flexbar (Dodt et al., 2012; Renaud et al., 2015) and deML (Renaud et al., 2015). Operational taxonomic units OTUs were identified and assigned to taxonomy from demultiplexed samples using the MJOLNIR pipeline (https://github.com/uit-metabarcoding/MJOLNIR) as in (Turon et al., 2022). MJOLNIR implements a variety of programs from the OBITOOLS package (Boyer et al., 2016) for sequence preprocessing and taxonomic assignment as well as VSEARCH (Rognes et al., 2016) and SWARM (Mahé et al., 2022) for chimera detection and OTU clustering respectively. The median read depth per sample was 220,000. Fourteen negative extraction controls were amplified and sequenced. Appreciable contamination (c.20K reads) was noted only in one. OTU taxonomic identities were passed directly to correlation analyses with biotic and abiotic variables, as well as with microscopy data.

### Data analysis

Plankton taxa identified by eDNA metabarcoding were aggregated at the genus level, unless there was only one species representing the genus in which case the name of the species was retained (e.g. *Lizzia blondina*). In microscopy data, the name of the genus was used when higher resolution could not be achieved with microscopy (e.g. *Chaetoceros*, *Pseudo-nitzschia*). We also created higher taxonomic level aggregations, eg phyla or classes depending on the purposes of the analysis and data visualisation. Metabarcoding data were transformed to relative reads by dividing the reads of each OTU by the total abundance in the sample. After transformation, we excluded from the OTU-read dataset all barcodes that corresponded to fish, birds, mammals and terrestrial plants. For certain analyses, we filled gaps in the daily samples in our timeseries using linear interpolation that excluded gaps at the edges of the dataset.

For multivariate analysis of microscopy and eDNA metabarcoding species data, we determined between-sample similarities using the Bray-Curtis index and visualized these with multidimensional scaling ordination using the “vegan” R package (Oksanen & et.al., 2022). For this analysis, we used non-interpolated and non-transformed species-abundance or OTU-reads data, aggregated at the genus level.

For modelling purposes, gaps along the timeline of daily data were filled using linear interpolation, excluding gaps at the edges of the timeline. To test for potential positive associations of plankton-borne organisms with the incidence of fish PGD and mortality we used a three-step conservative model selection process of plankton predictors. As a first step, for each of the two sites separately, we selected all species that showed a >0.4 Spearman correlation with either PGD or mortality. As a second step, we fitted a linear model with PGD or mortality as response variables and as explanatory variables the highly correlated species that were selected from step 1. In addition, here we also accounted for temperature and oxygen as covariates in the model due to the direct effect they can exert on fish health. Specifically, dissolved oxygen concentration is a known predictor of fish welfare (Remen et al., 2016) and more recently temperature has also been reported as a direct predictor of gill pathology (Herrero et al., 2022) and has been found to alter the bacterial microbiome in fish gills associated with disease (Ghosh et al., 2022). Therefore, to get the individual effect of each species, after accounting for temperature, oxygen and the other species present, we used the *glmm.hp* algorithm and corresponding R package (Lai et al., 2022) to fit two models with PGD and mortality, each including temperature, oxygen and the selected correlated species as explanatory covariates. The *glmm.hp* algorithm determines the relative importance of collinear predictors by partitioning variation explained by each covariate into unique and average shared and is recommended for datasets with multiple collinear variables such as ours. The sum of these two components (unique and average shared variation) was used to express the percentage of the individual contribution of each predictor species in our model. The selected species from step 1 were added to the model with forward selection starting from the highly correlated and proceeding to the weakly correlated while discarding those with <3% individual effect on the total variation in either PGD or mortality. As a third step, we determined whether the effect of the selected species depended on the site exposure level. To achieve this, we merged the data from both sites and fitted a PGD and a mortality model using as explanatory terms the temperature, oxygen, and the factor exposure level (exposed/sheltered) as well as all the species selected from step 2 and their interaction with exposure level. All explanatory covariates were scaled for inclusion in the models. This three-step process was then repeated to test the effect of lagged PGD and mortality data by 2, 5 and 10 days behind the plankton species data to establish any lag effects on fish health. The lagging of data was performed on the fish condition data using the package “lubridate” (Grolemund & Wickham, 2011) and subsequently, this lagged dataset was merged with the non-lagged species-abundance dataset. All analysis was carried out in R v.4.3.1. (R Core Team, 2022).

## RESULTS

Fish mortality within each of the two aquaculture sites presented strong temporal variability with notable increases observed from June onwards. In the exposed site, mortality peaked in late June and September whereas in the sheltered site, the highest mortalities were recorded in August and September 2021 (Figure 2). PGD scores also peaked in late summer and were overall higher in the sheltered site, where less variability was observed. Total abundance of phytoplankton and zooplankton presented multiple peaks throughout the study period, with higher abundances during the summer months particularly in the sheltered site.

**Figure 1.**
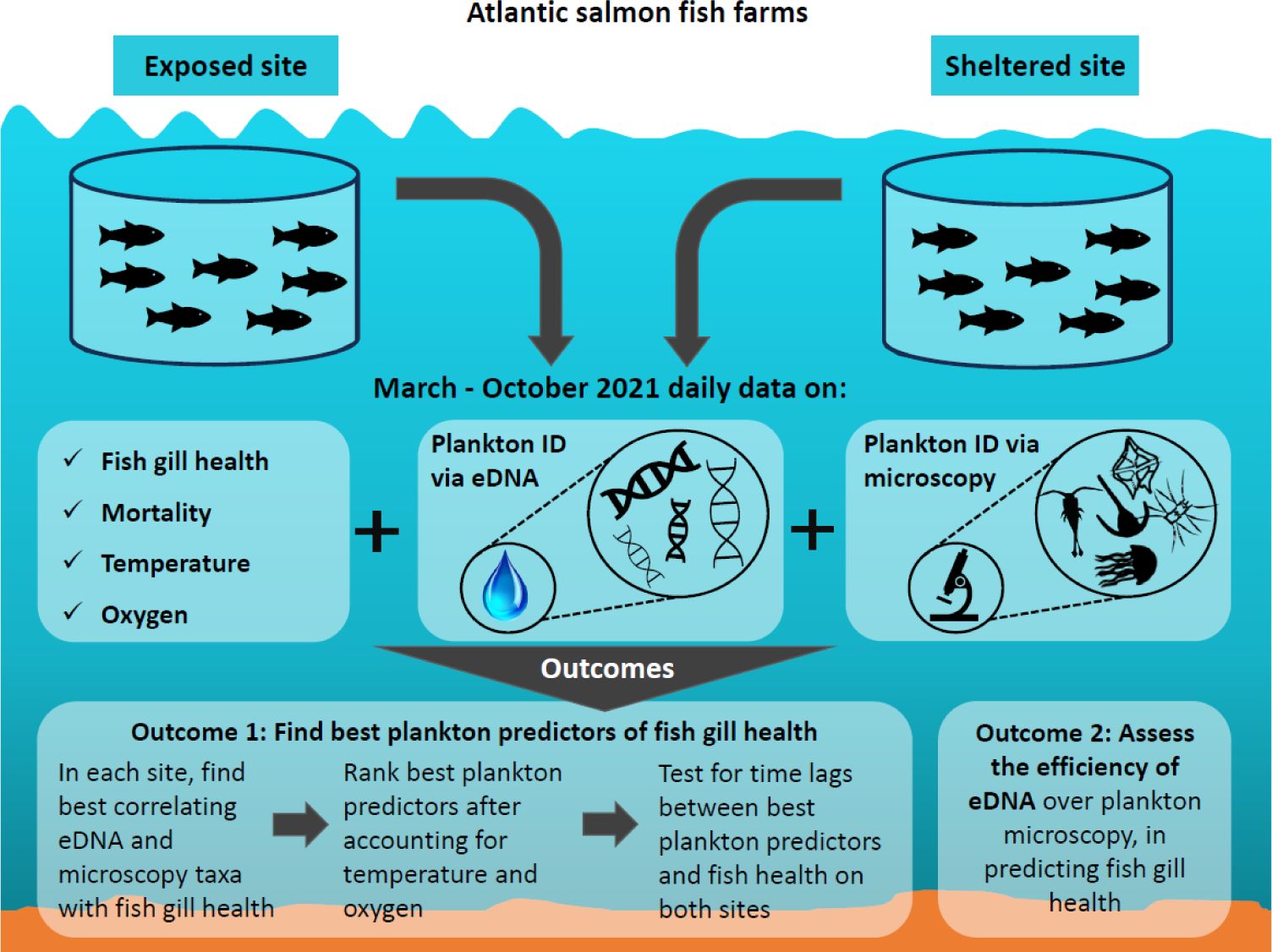
Approach, outputs, and applicability of the methodological framework of our study. Plankton symbol attribution: ian.umces.edu/media-library

**Figure 2.**
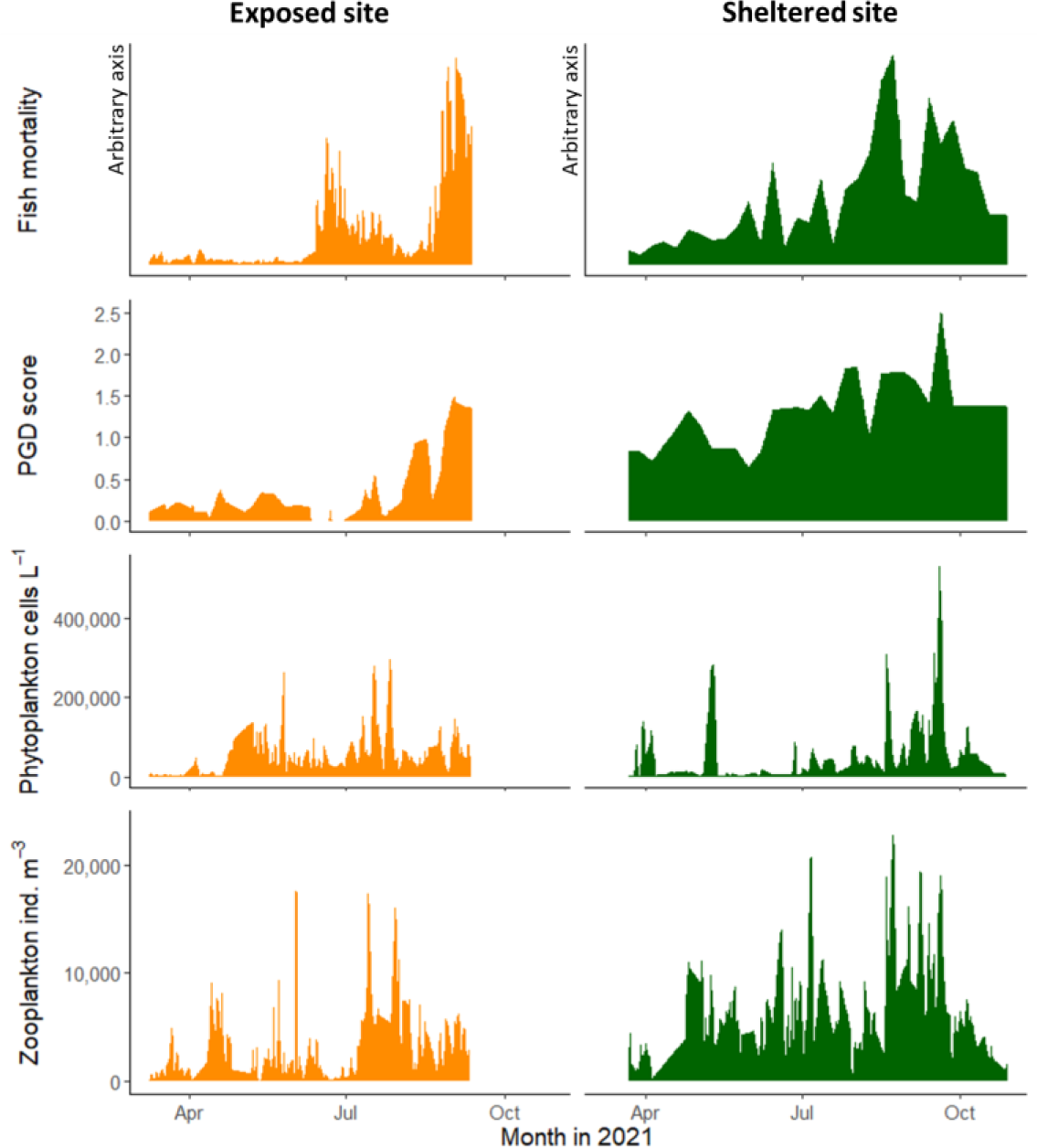
Dynamics of fish condition variables and total phytoplankton and zooplankton abundance. -via microscopy-during the study period (March to October 2021). Trends are shown for an exposed to the open sea versus a sheltered aquaculture site. Fish mortality is based on z-scores corrected for negative values and is not directly comparable between the two sites.

The relative abundance (as relative reads in the sample) of dominant phyla identified by eDNA metabarcoding presented temporal and geographical differences (Figure 3A). The exposed site was characterized by a large number of unassigned OTUs, and the top 6 phyla in decreasing abundance were Unassigned, Discosea, Ascomycota, Bacillariophyta, Cnidaria and Rotifera (Figure 3B). For the sheltered site, the top 6 phyla in decreasing abundance were Discosea, Unassigned, Rotifera, Bacillariophyta, Cnidaria and Ascomycota.

**Figure 3.**
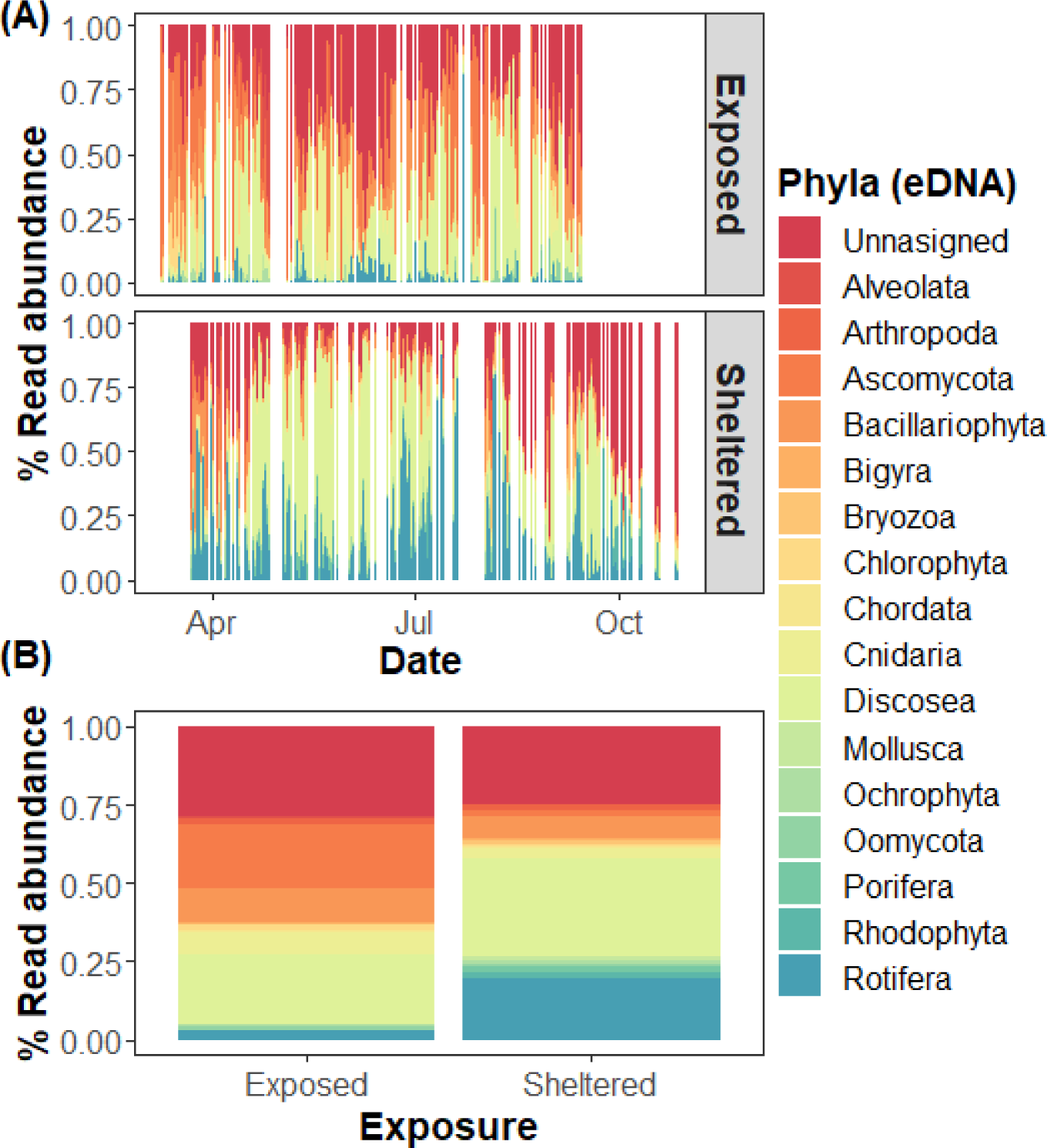
Classification of relative abundances of Operational Taxonomic Units (OTUs) into the most abundant phyla identified via eDNA metabarcoding across our sampling period (March to October 2021) and two aquaculture sites (exposed and sheltered). Taxonomic identities of OTUs and thus Phyla were manually curated using BLASTn and only the Phyla contributing to >1.5% of reads were included in the graph. OTUs with no known taxonomic classification are shown as ‘Unassigned’.

Plankton species composition was different between the exposed and sheltered sites (Figure 4A), and the difference was more distinct when eDNA metabarcoding data were used (Figure 4C). A seasonal pattern in species composition was also observed on both sites with the spring species composition being different to the late composition (Figure 4B,D). This was due to the fact that most plankton taxa such as *Rhizosolenia setigera*, *Chaetoceros*, *Dictyocha*, *Lizzia*, and *Oikopleura*, showed increased abundances from June onwards. Very few taxa showed increased abundance throughout the production period such as the Copepods and the diatom *Pseudo-nitzschia* (see also Figure 6).

**Figure 4.**
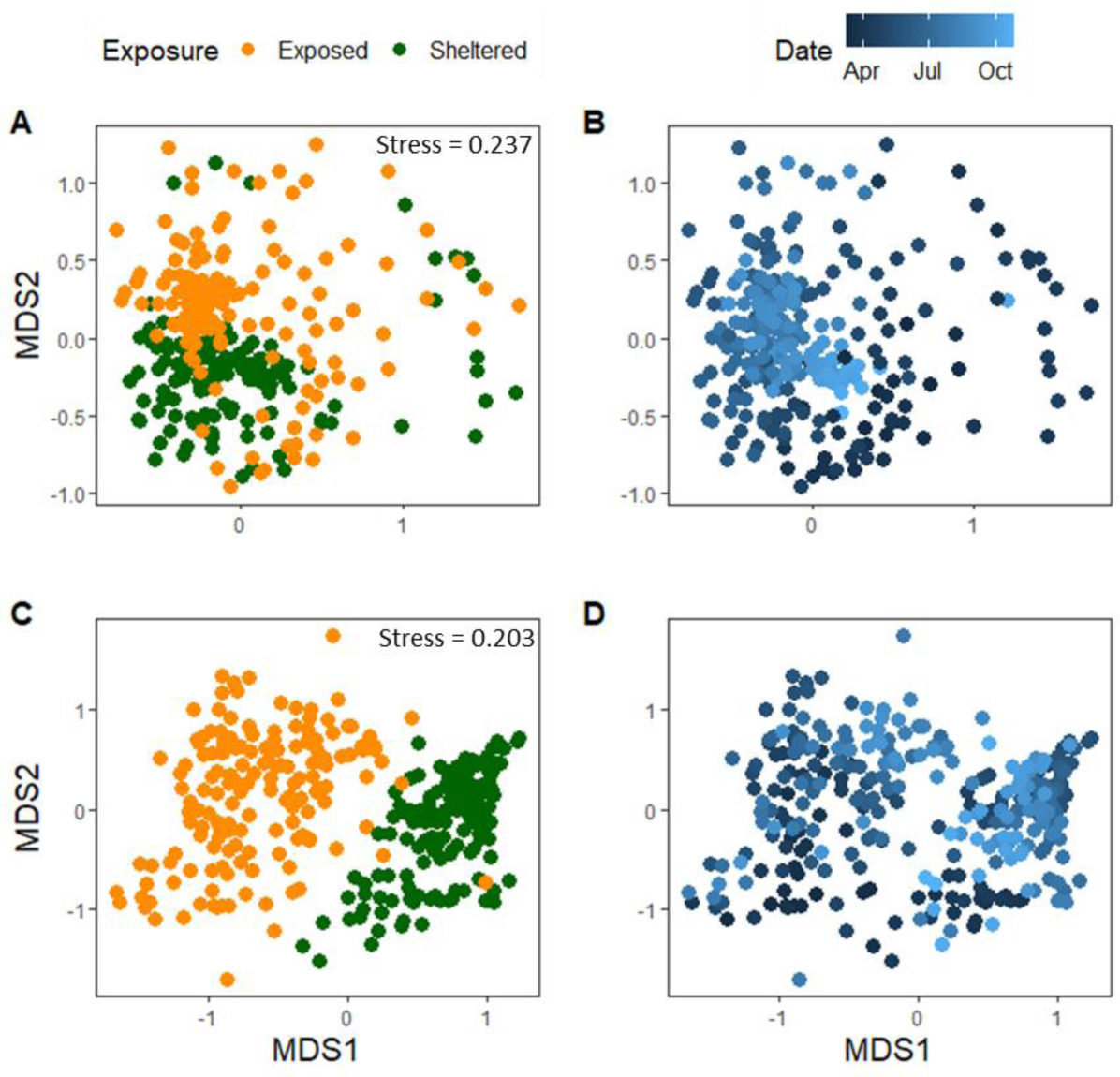
Multi-dimensional scaling ordination showing the relative similarity between our total number of 412 samples of species-abundances as identified by microscopy (panels A and B) and eDNA metabarcoding (panels C and D). Analysis was based on the Bray-Curtis similarity index calculated on non-interpolated, species-abundance microscopy data and genus/relative read metabarcoding data.

**Figure 5.**
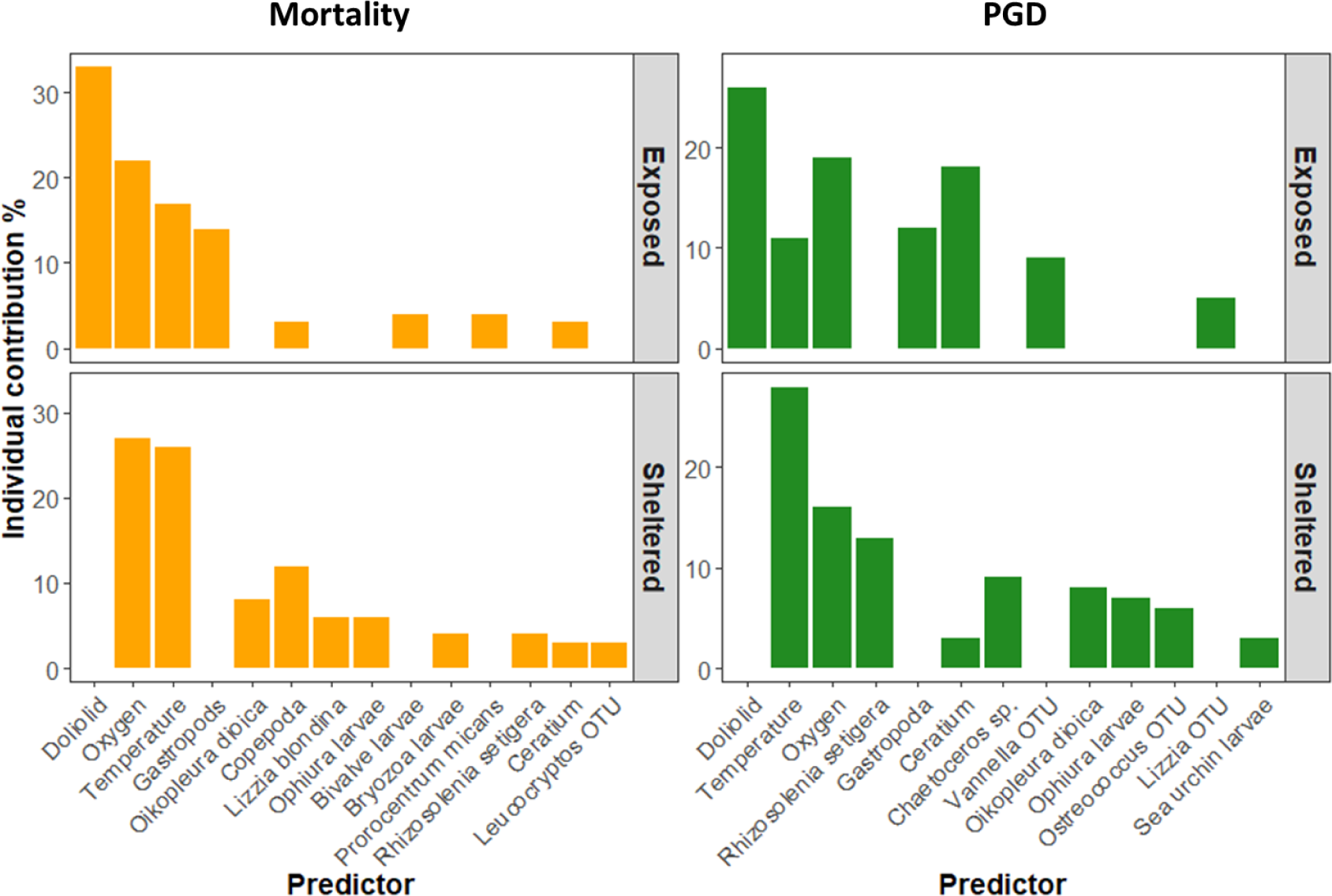
Ranking of the relative importance of predictor variables for fish PGD and mortality within sites. The ranking is conservative and is based on a preselection of predictors whereby those correlated with Spearman R2<0.3 with either PGD or mortality were excluded. The model included the top predictors which have an individual contribution of >3% to the overall shared variation in PGD and mortality.

**Figure 6.**
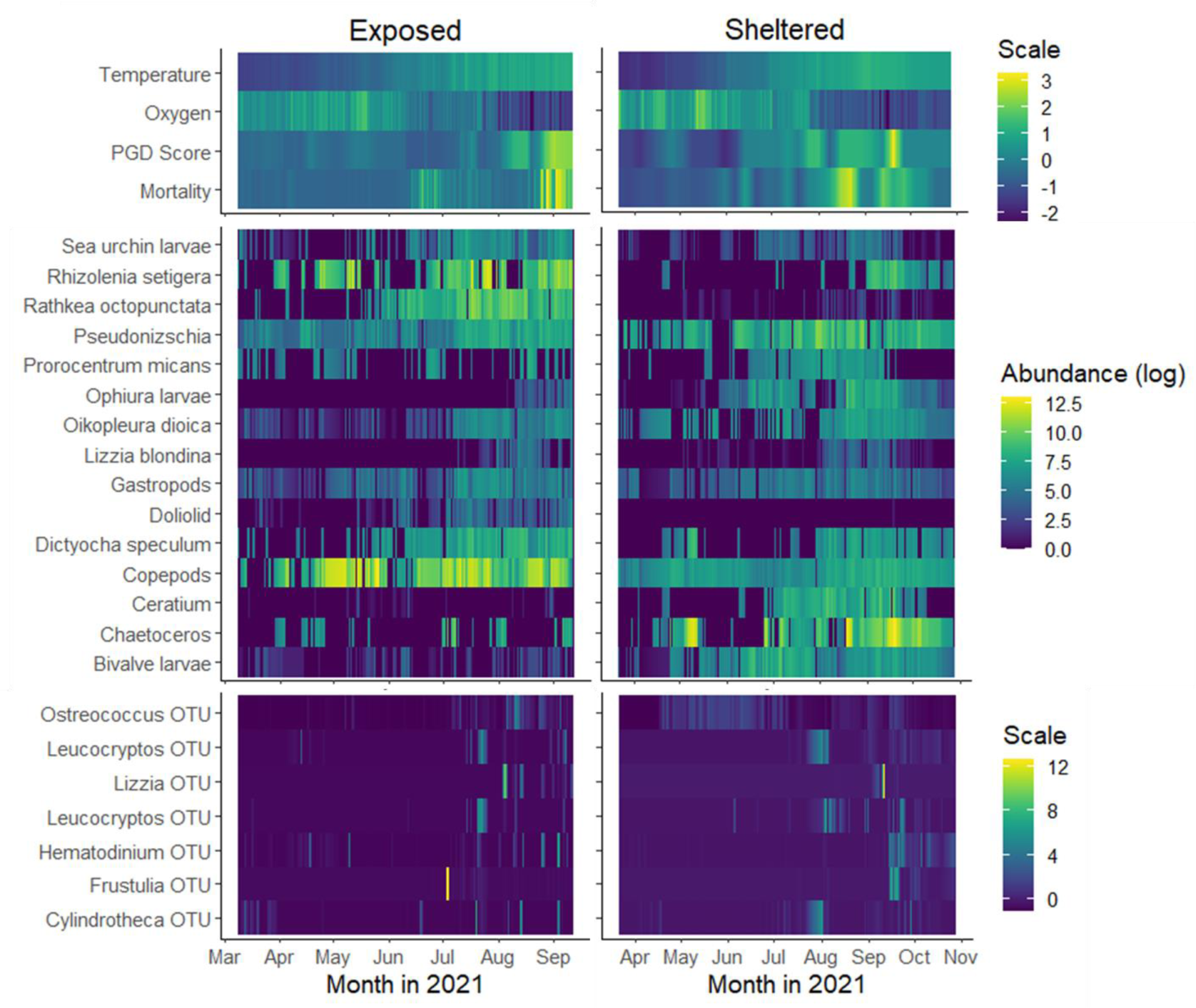
Heatmaps showing the dynamics of variables related to the environment, fish health and plankton-borne vectors. Plankton was identified either morphologically, or molecularly (OTU suffix) and those shown are the good predictors of PGD and mortality in at least one of the two sites (exposed and sheltered). All variables have been interpolated and genus OTUs, fish condition, and abiotic variables have been normalised by zero-centering to enable direct comparisons with microscopy data. Abundance was measured in cells/L or individuals/m^3^ depending on whether the organism was a planktonic unicellular eukaryote (protist) or zooplankton.

Temperature increase and dissolved oxygen decrease were significantly and strongly associated with fish mortality and PGD prevalence at both sites (Figure 5). Multiple phytoplankton and zooplankton taxa were positively associated with PGD and mortality and this effect was often dependent on the site. For example, doliolids were only present in the exposed site, where they appeared to be associated with both mortality and PGD whereas the appendicularian *Oikopleura dioica* was present on both sites and its effect on PGD was site-dependent (Table 1).

**Table 1.** Plankton taxa positively associated with the incidence of fish PGD and mortality. . Results are shown when no lag was assumed between fish health and plankton abundances and for lagging the PGD and mortality data by 2, 5 and 10 days behind the environment and plankton species data. A cross indicates a significant positive association of the species with either PGD or mortality after accounting for the effect of temperature and oxygen saturation. A grey highlight shows that the effect depended on the site/exposure level. The OTU label indicates where the taxon was detected via molecular means.

| Taxon | PGD |  |  |  | Mortality |  |  |  |
| --- | --- | --- | --- | --- | --- | --- | --- | --- |
|  | 0d | 2d | 5d | 10d | 0d | 2d | 5d | 10d |
| *Attheya* OTU |  |  |  |  |  | x | x | x |
| *Ceratium* | x | x |  | x | x | x | x | x |
| *Chaetoceros* sp | x | x |  |  |  |  |  |  |
| *Cylindrotheca* OTU |  |  | x | x |  |  |  |  |
| *Dictyocha speculum* |  |  |  | x |  |  |  |  |
| *Hematodinium* OTU |  |  | x |  |  |  |  | x |
| *Leucocryptos* OTU |  | x |  |  | x |  | x |  |
| *Ostreococcus* OTU | x | x | x | x |  |  |  |  |
| *Prorocentrum micans* |  |  |  |  | x | x |  | x |
| *Pseudo-nitzschia* |  |  |  |  |  |  | x | x |
| *Frustulia* OTU |  |  |  |  |  |  | x |  |
| *Rhizosolenia setigera* | x | x |  |  | x | x | x |  |
| Bivalvia larvae |  |  |  |  | x | x |  | x |
| Copepods | x | x | x | x | x | x | x | x |
| Doliolid | x | x | x | x | x | x | x |  |
| Gastropoda | x | x |  |  | x | x | x | x |
| *Lizzia blondina* |  |  |  | x | x |  |  |  |
| *Lizzia blondina* OTU | x | x |  |  |  |  |  |  |
| *Oikopleura dioica* | x | x | x | x | x | x | x | x |
| Ophiura larvae | x | x |  |  | x | x | x | x |
| *Rathkea octopunctata* |  |  |  |  |  | x | x |  |
| Sea urchin larvae | x | x |  |  |  |  |  |  |
| *Vannella* OTU | x | x |  |  |  |  | x |  |

*Ceratium*, doliolid, *Oikopleura* and *Ophiura* larvae were associated with both PGD and mortality, whereas some species were uniquely associated with either PGD or mortality. Specifically, *Chaetoceros*, *Ostreococcus*, *Cylindrotheca* and sea urchin larvae were uniquely associated with PGD, whereas *Attheya*, *Prorocentrum micans*, *Pseudo-nitzschia*, *Frustulia* and *Rathkea octopunctata* were uniquely associated with fish mortality (Table 1). Some plankton species appeared associated with PGD and mortality only when lags were included at 2, 5 or 10 days, reflecting a delayed impact. Specifically, *Cylindrotheca* was associated with PGD with a 5- and a 10-day lag (Table 1) and *Attheya*, *Pseudo-nitzschia* and *Rathkea* were associated with mortality when the later was lagged by more than 2 days.

The abundance of plankton species which showed a significant association with fish mortality and PGD showed strong temporal variation throughout the study period from March to October 2021 with most species increasing in abundance over the summer months (Figure 6). An exception to this summer increase were copepod, gastropod larvae and bivalve larvae, whose abundance was high throughout the period on both sites and diatom species such as *Rhizosolenia* and *Chaetoceros* which also showed spring peaks in abundance. Plankton abundances based on the microscopy showed more pronounced temporal autocorrelation patterns than eDNA metabarcoding (see taxa OTU taxa in Figure 6).

At genus level, abundant taxa that were identified by microscopy but not as OTUs and vice versa, were also observed. An example of this was the zooplankton species *O. dioica*, where no corresponding OTU was observed from the molecular data. On the other hand, eDNA metabarcoding identified the diatom species *Cylindrotheca* as well as several amoebozoan genera which was not detected in the plankton samples via microscopy. For the species that were identified by both methods, not all genera showed significant correlations between abundances (microscopy) and reads (metabarcoding) (Figure 7).

**Figure 7.**
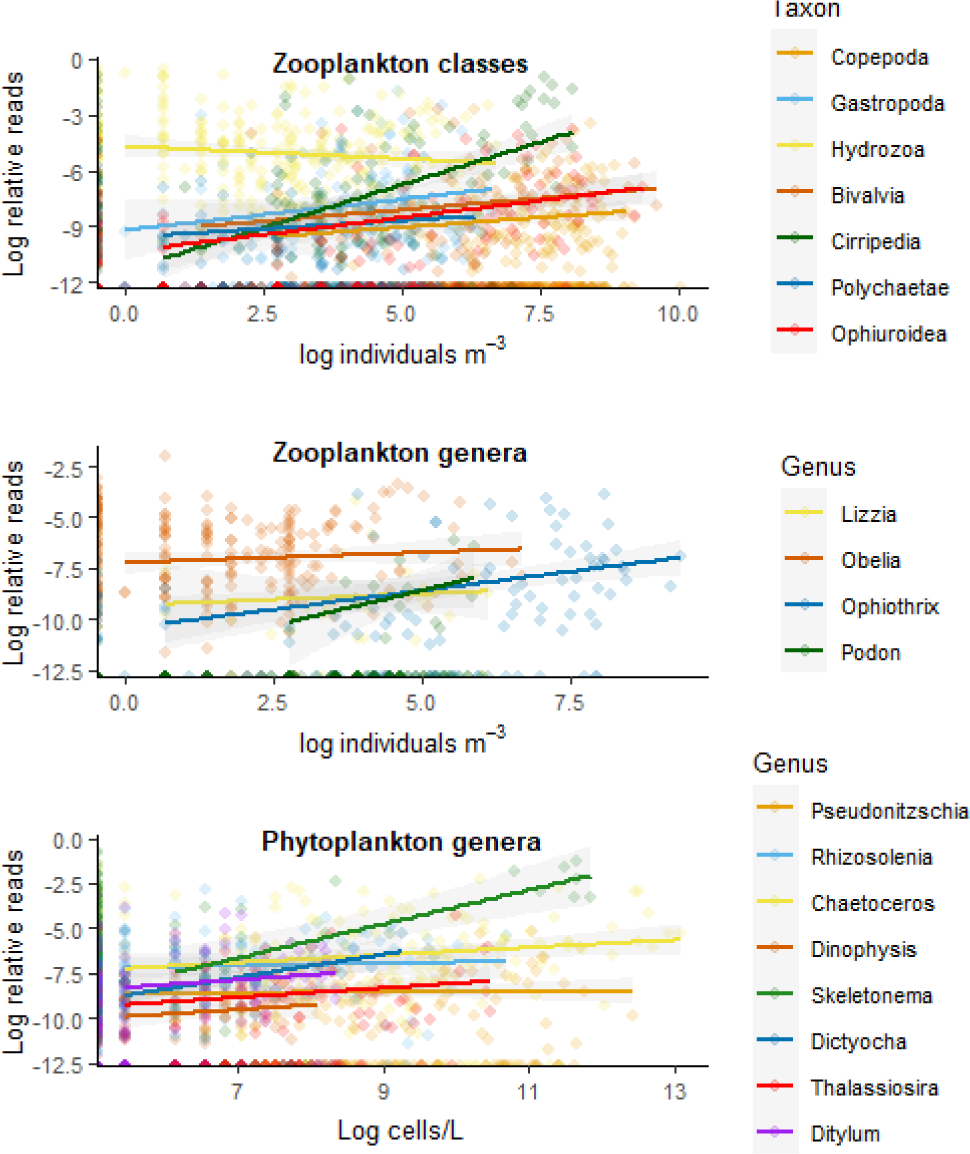
Relationship between abundances of classes and genera as identified by eDNA metabarcoding. (log relative reads) and microscopy (log cells/L or Ind/m3). Site specific relationships between the two methods are shown in the supplementary material (S1) and statistical results presented in Table 2.

To further assess congruence between metabarcoding and microscopy in our dataset, we evaluated the relative sensitivity of eDNA and microscopic data for taxa that were clearly identified by both methods on any given day (Table 2). We noted a high level of discrepancy in detection sensitivity (a taxon was identified by one technique, but not the other). For example, over 412 sampling days, spanning both locations, at best *Skeletonema* detection discrepancies were detected in 42% of the samples and worst *Ditylum* in 82% of the samples (Table 2). For those taxa identified by both techniques, the relative sensitivity of each was evaluated as the ratio of respective detection days, which revealed that for some taxa, such as the diatom *Skeletonema*, eDNA was more sensitive while for others, such as the hydrozoan *Lizzia,* microscopy performed better.

**Table 2.**
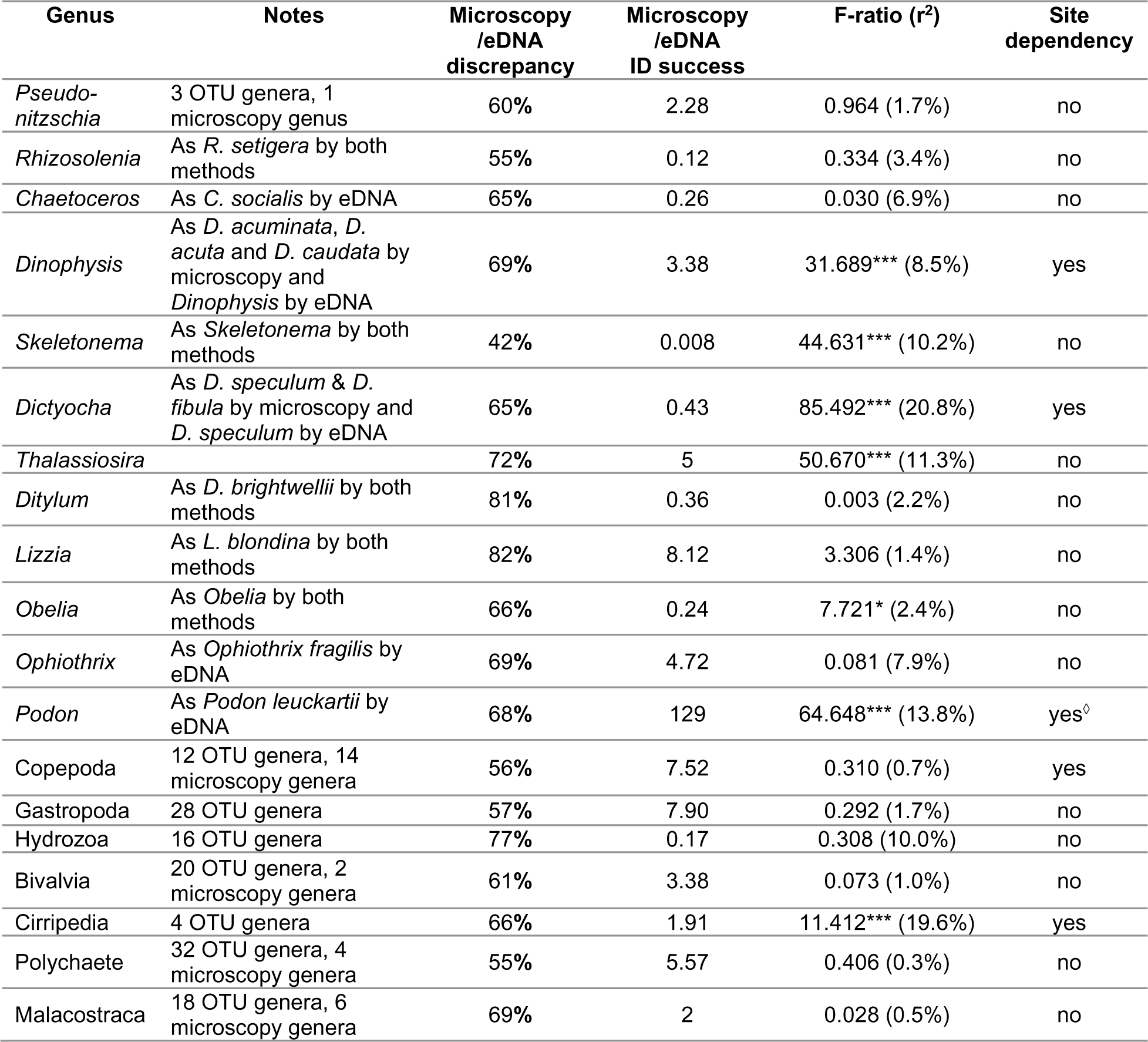
Assessment of congruence between morphological and eDNA metabarcoding plankton data across those genera and broader taxonomic groups that were identified by both methods (e.g. Appendicularia were not detected by eDNA and are thus excluded). The degree of discrepancy between the two methods is expressed as the percentage of total samples (412) that a genus was identified by only one of the two methods. The ratio of samples identified by microscopy over those identified by eDNA shows the relative strength of each method (>1 indicates that microscopy is more effective whereas <1 indicates eDNA is more effective in identifying this species). The strength of relationship between eDNA-relative read data and abundance microscopy data is assessed using linear regression that assumes dependency on the exposure level/site (Genus_OTU_∼Genus_microscopy_ x Exposure level). The strength of the relationship is expressed by the F-ratio, the significance level (*** significance at 99.9% level, ** at 99%, * at 95% and no asterisk indicates no significance) and the r-squared.^◊^ the taxon only present on the exposed site.

| Genus | Notes | Microscopy<br>/eDNA<br>discrepancy | Microscopy<br>/eDNA<br>ID success | F-ratio (r<sup>2</sup>) | Site<br>dependency |
| --- | --- | --- | --- | --- | --- |
| *Pseudo-nitzschia* | 3 OTU genera, 1 microscopy genus | 60% | 2.28 | 0.964 (1.7%) | no |
| *Rhizosolenia* | As *R. setigera* by both methods | 55% | 0.12 | 0.334 (3.4%) | no |
| *Chaetoceros* | As *C. socialis* by eDNA | 65% | 0.26 | 0.030 (6.9%) | no |
| *Dinophysis* | As *D. acuminata*, *D. acuta* and *D. caudata* by microscopy and *Dinophysis* by eDNA | 69% | 3.38 | 31.689*** (8.5%) | yes |
| *Skeletonema* | As *Skeletonema* by both methods | 42% | 0.008 | 44.631*** (10.2%) | no |
| *Dictyocha* | As *D. speculum* & *D. fibula* by microscopy and *D. speculum* by eDNA | 65% | 0.43 | 85.492*** (20.8%) | yes |
| *Thalassiosira* |  | 72% | 5 | 50.670*** (11.3%) | no |
| *Ditylum* | As *D. brightwellii* by both methods | 81% | 0.36 | 0.003 (2.2%) | no |
| *Lizzia* | As *L. blondina* by both methods | 82% | 8.12 | 3.306 (1.4%) | no |
| *Obelia* | As *Obelia* by both methods | 66% | 0.24 | 7.721* (2.4%) | no |
| *Ophiothrix* | As *Ophiothrix fragilis* by eDNA | 69% | 4.72 | 0.081 (7.9%) | no |
| *Podon* | As *Podon leuckartii* by eDNA | 68% | 129 | 64.648*** (13.8%) | yes<sup>o</sup> |
| Copepoda | 12 OTU genera, 14 microscopy genera | 56% | 7.52 | 0.310 (0.7%) | yes |
| Gastropoda | 28 OTU genera | 57% | 7.90 | 0.292 (1.7%) | no |
| Hydrozoa | 16 OTU genera | 77% | 0.17 | 0.308 (10.0%) | no |
| Bivalvia | 20 OTU genera, 2 microscopy genera | 61% | 3.38 | 0.073 (1.0%) | no |
| Cirripedia | 4 OTU genera | 66% | 1.91 | 11.412*** (19.6%) | yes |
| Polychaete | 32 OTU genera, 4 microscopy genera | 55% | 5.57 | 0.406 (0.3%) | no |
| Malacostraca | 18 OTU genera, 6 microscopy genera | 69% | 2 | 0.028 (0.5%) | no |

## DISCUSSION

Our survey represents a thorough assessment of the planktonic threats faced by an open-water marine aquaculture species, both in terms of breadth (the number of planktonic taxa identified) and depth (number and temporal frequency of sampling days). We noted a general trend in fish health for the two sites we monitored, with mortality and PGD scores worsening in late summer months (July-September). High sampling frequency and two sites with contrasting environmental characteristics enabled some statistical deconvolution of planktonic species associated with poor fish health from collinear variables such as sea surface temperature and dissolved oxygen. Biological correlates with PGD and mortality were largely site-specific. As such, only the dinoflagellate genus *Ceratium,* copepod larvae and the hydromedusa *Lizzia blondina* were linked with poor fish health at both sites. Meanwhile, doliolids (classified as *Doliolum nationalis* and *Dolioletta gegenbauri*) were strongly associated with poor fish health at the exposed site, and the appendicularian *O*. dioica was a principal biological correlate with mortality at the sheltered site. Interestingly, the correlation between some planktonic taxa (e.g. *Pseudo-nitzschia*, *Cylindrotheca*) and poor fish health was noted only when fish health variables were lagged behind planktonic dynamics, indicating delayed effects of certain planktonic species on fish condition. Finally, we evaluated the correspondence between molecular (eDNA metabarcoding) and morphological (microscopy) approaches in plankton detection. In general, the approaches were poorly correlated, with sequence read abundances being a poor predictor of microscopic cell / organismal abundance. Relative detection sensitivities were also highly variable, with some important planktonic predictors of fish health being detected by only one of the two techniques deployed (microscopy vs eDNA).

The summer months have represented a growing challenge to aquaculture producers over recent years in terms of gill health. Sporadic reports of seasonal gill challenges date back almost twenty years (e.g. Boerlage et al., 2020). On the west coast of Ireland, a southerly sentinel for salmonid aquaculture in the Northern Hemisphere, high gill-related mortalities from June to September have been reported for over a decade (Rodger et al., 2011). It is unclear to what extent these gill challenges follow warming sea temperatures, but the summer trend for gill deterioration during summertime in Scottish waters now seems established. Salmon Scotland reports late summer mortalities increasing steadily year on year, e.g. September 2018 - 1.57%, September 2022 - 4.65% (Salmon Scotland, 2022). Our data are consistent with this trend but also point to additional peaks in mortality earlier in the summer. The current increasing trend in fish mortalities in addition to sea temperature rise and increasing plankton blooms raise concerns for the future of the sector. Understanding the aetiology of fish disease is therefore essential in order to develop and implement mitigation strategies.

Temperature and dissolved oxygen have been reported as direct drivers of salmon gill health (Ghosh et al., 2022; Herrero et al., 2022; Remen et al., 2016) and our study confirms this. However, having accounted for the effect of these key abiotic drivers, our analyses of the data across both sites identified numerous biological correlates with poor fish gill health and mortality. Cnidarian gelatinous zooplankton, especially hydrozoans, have been widely reported as drivers of gill insult and salmon mortality (Boerlage et al., 2020). Our morphological analyses detected multiple hydrozoan species – *Rathkea octopunctata, Lizzia blondina* (Figure 6), *Obelia* spp.*, Ectopleura* sp.*, Bougainvillia* sp.*, Phialella quadrata, Clytia hemisphaerica* and others (Supplementary data). The hydroid of *Ectopleura* cf. *larynx*, a cnidarian fouling species with the potential to cause salmon health issues (Bloecher et al., 2018) was identified from our zooplankton samples, presumably released into the water during regular in-situ net cleaning in the farms. Only *Rathkea, Lizzia* and *Obelia* were detected in appreciable abundance, and only *L. blondina* correlated with gill damage (PGD, Exposed Site) and mortality (Sheltered site). *Rathkea* also had some impact on mortality but the impact lagged by several days after detection of the species. *L. blondina* has been correlated with gill damage in Scotland previously, as have *Obelia* sp. (Kintner & Brierley, 2019). Despite the high abundance of *Obelia* across both sites, much higher than *L. blondina* (Supplementary data), we did not identify any correlation with fish health. Other hydrozoan species such as the siphonophore *Muggiaea atlantica*, for example (e.g. Baxter et al., 2011) are known to represent particular threats to salmon gills although they were not detected in our study. The pathophysiology of the different salmon-hydrozoan species-specific interactions might be determined by the mechanical damage of the nematocysts as well as the toxins released (see Bloecher et al. 2018), however other factors such as jellyfish swarm size, exposure time (Clinton et al., 2021), toxins type and fish immune response could also have an effect. Moreover, our results show that gelatinous zooplankton (GZP) causing salmon gill disease include not only cnidarians but other groups such as appendicularians and doliolids —which do not have nematocysts to penetrate the gill tissue— suggest that jellyfish envenomation might not the only trigger of CGD, and certainly calls for further investigation on the mechanisms that determine the pathogenicity of this group.

Diatom blooms have long been understood as drivers of salmon mortality at sea (Albright et al., 1993; Bell, 1943). Here, multiple diatoms (e.g. *Rhizosolenia*, *Chaetoceros, Pseudo-nitzschia*) were correlated with PGD and/or mortality. Proposed mechanisms include both mechanical damage of gill tissue by the siliceous spines of diatoms (Albright et al., 1993) as well as direct toxin production by microalgae species (e.g. Bates et al., 2018). The dynamics of dinoflagellates *Ceratium* and *Prorocentrum* were also negatively associated with fish health. Bloom concentrations of *Ceratium* and *Prorocentrum* have been linked with dissolved oxygen depletion and subsequent fish kills in the past (Azanza et al., 2005; Glibert et al., 2002; Malone, 1978). In our study, *Ceratium* and *Prorocentrum* reached maximum concentrations of 3.5x10^4^ and 7x10^3^ cells/L respectively, therefore, although high, neither is considered bloom concentration. However, *Ceratium* consisted of species such as *C. lineatum*, *C. furca*, *C. fusus* and *C. tripos,* the later three being quite voluminous dinoflagellates shapes as needles or u-shaped with multiple long horns that can reach up to 230 μm in size. Lower concentrations of large *Ceratium* species have been associated to fish kills in the past via mechanical damage to fish gills and secondary infection (Orellana-Cepeda et al., 2002) which might be the also the case in our study. Finally, the silicoflagellate, *Dictyocha speculum* also had a lagged effect on the incidence of PGD in our study, in agreement with previous report on this microalgae causing mass mortality on farmed *S. salar* in the Galician coastal waters (Prego et al., 2023).

Among the most highly ranked taxa in terms of their correlation with both PGD and salmon mortality were several gelatinous zooplankton species, as well as unicellular eukaryotes, never before suggested as harmful to fish. These species provide a first glimpse below the tip of the iceberg in terms of the hidden diversity of planktonic threats to gill health. Doliolids are planktonic, filter-feeding tunicates 1-2 mm in length, which appeared strongly associated with mortality and PGD at the exposed site, more so than temperature and DO (Figure 5). Doliolid blooms have been associated with climate-change related heatwaves (Pinchuk et al., 2021) and these numbers broadly tracked temperature in our study. Doliolids are not known to be toxic, although their selectivity for diatoms and ciliates (Frischer et al., 2021) suggest that toxins accumulation might be occurring in doliolids bodies; this, however, has not been tested to date, to the best of our knowledge. Alternatively, these small GZP could simply adhere to and clog the gill lamellae, obstructing gaseous exchange. Another pelagic tunicate strongly associated with gill disease and mortality, this time at the sheltered site, was the appendicularian *O. dioica*. Again, the role of *O. dioica* as an irritant is far from clear. This species filter feeds on microplankton, including marine viruses, via an extruded cellulose net (Lawrence et al., 2018). *O. dioica* may potentially bioaccumulate heavy metals, or even toxins of algal origin, however, they too can be highly sensitive to such compounds (Calatayud et al., 2018; Torres-Águila et al., 2018). As with doliolids, a mechanism of *O. dioica* toxicity beyond direct obstruction of the gas exchange surface has not been defined yet.

As well as doliolids and appendicularians, several other components of the zooplankton community such as sea urchin larvae, copepods and gastropods have been identified as potential poor gill health and mortality drivers. As this study initially targeted planktonic groups causing diminished salmon health based on literature (i.e. cnidarians and harmful phytoplankton blooms), morphological identification efforts did not focus at lower taxonomic levels on these common and highly diverse taxa. At the other end of the scale, the amoebozoan *Vanella* and the protist *Leucocryptos* were also incriminated. The correlations we detected, however, suggest that zooplankton-gill health interactions can involve more players and levels of complexity than traditionally expected. A mechanism that might link each class of organism to gill damage is beyond the reach of this study but is clearly an important avenue for further research. However, it is worth noting that the most important correlates with poor gill health and mortality were not limited to hydrozoans or harmful algal species, despite the fact that these groups are the only ones reported in the literature (e.g. Boerlage et al., 2020), and the species most closely monitored by the aquaculture industry (Bickerdike, pers comm). Indeed, the most important correlates in this study have never before been reported as important drivers of gill health or mortality of farmed Atlantic salmon.

The presence of a planktonic irritant on a given day may not result in an immediate impact on either gill integrity or fish health. Furthermore, gill damage is likely to be cumulative (Boerlage et al., 2020; Østevik et al., 2022) with the condition worsening over the course of spring/summer months, increasing the sensitivity of fish to mortality through physiological stress associated with feeding, medicine treatments and other handling events. Cumulative damage may be best reflected through PGD score, as AGD scores do seem to improve after freshwater treatments for *N. perurans* (e.g. Parsons et al., 2001). The ability of AGD gill lesions to resolve post-treatment is clearly evidenced in our data, whereby high AGD gill scores, which trigger treatment events, has a strong negative relationship with mortality independently of site (F-value=91, p<0.001). In contrast, PGD score was positively associated with mortality independently of site (F-value=615, p<0.001). To capture some of the delayed and/or cumulative impacts of different irritants on gill health, we lagged mortality and PGD behind our planktonic data and assessed how strongly different taxa were associated with poor health outcomes. Several taxa identified in our unlagged ranking analysis also appeared to have significant lagged associations with mortality. However, several previously unrecorded phytoplankton taxa may have a delayed, but nevertheless important, impact on fish health, e.g. the diatoms *Cylindrotheca* and *Frustrulia*, the Ochrophyte *Dictyocha* and the dinoflagellate *Hematodinium*. *Hematodinium* is of particular interest as it is a parasite of a decapod crustacean of economic importance in Scotland and globally (Beevers et al., 2012), although to our knowledge it has never been isolated from Atlantic salmon.

Establishing the value of eDNA in predicting organismal biomass is a long-standing goal of molecular ecological research in freshwater and marine environments (e.g. Bourque et al., 2022; Lamb et al., 2019; Rourke et al., 2022). Experimental work has demonstrated that eDNA concentrations tracked via qPCR (e.g. Bourque et al., 2022) and metabarcoding (Peters et al., 2018) can predict absolute and/or fold changes in plankton biomass. Some success was also reported in the marine environment in field conditions (Ershova et al., 2021; Santi et al., 2021). In our study, results were mixed. For zooplankton taxa, the predictive value of eDNA on biomass was generally poor with the exception of Cirripedia which showed the strongest relationship between microscopy and metabarcoding data with 20% of the variation explained by the linear relationship. There is a stronger relationship between molecular and microscopic estimates of abundance for phytoplankton, especially for more frequently occurring taxa (e.g. *Skeletonema*). The higher biomass disparity within and among zooplankton taxa might have contributed to this result, and experimental validation of the relationship between eDNA detection efficiency and biomass for individual zooplankton species may complement our work. The sensitivity of eDNA and microscopic approaches also varied, with important taxa often clearly identified by one technique and largely missed by another. Examples of this were the diatom *Attheya* that was identified by metabarcoding and which can be confused with the genus *Chaetoceros* during microscopy analysis. Another example was the diatom *Cylindrotheca* which was identified by metabarcoding but was not seen in the microscope samples potentially due to long sample preservation times.

For studies that compare traditional and DNA-based approaches, the conclusion that best serves the end goal of surveying biodiversity is often ‘use both’ (e.g. Santi et al., 2021). In some respects, our study is no exception and no single approach has a clear advantage. However, several potential improvements, especially to the molecular methodologies, could be considered to improve the detection of species of interest. First, despite mining our amplicon data for doliolid and appendicularian species, it appears that the mlCOIintF-XT/jgHCO2198 primer pair does not amplify tunicate COI with appreciable efficiency. Furthermore, reference sequences in the Universal-databank for Fisheries and Aquaculture cytochrome oxidase I database (https://github.com/uit-metabarcoding/DUFA) for these groups are missing or is incomplete, and further curation may be required (*Præbel.*, pers comm). Second, although the aim of this study was to survey planktonic diversity by as unbiased a means as possible, it is clear that a number of key eukaryotic salmon pathogens are missing from the molecular data. AGD gill scores and gill qPCR data indicate that *N. perurans* is abundant at both sites (See supplementary data), and, as in previous work (Bridle et al., 2010), *N. perurans* should be readily detected from the water column. Similarly, sea louse counts on salmon (*L. salmonis* / *Caligus elongatus*) indicate the presence of these parasites at both sites. DNA from adult and juvenile lice should also be abundant in the water column (e.g. Krolicka et al., 2022). However, unambiguous *N. perurans* reads were extremely rare in the dataset (19 total, see supplementary data), as were those for *L. salmonis* (976 total) and *C. elongatus* (18 total). For context, we found 876 red deer (*Cervus elaphus*) reads in the data, presumably washed in from streams and rivers. Clearly ‘non-target’ species abound in the data. Billions of reads in our dataset, for example, were assigned to free living, apparently non-pathogenic amoeba (e.g. *Cunea, Parvamoeba*), and the read depths required to per sample to ‘sequence through’ this biological noise using universal metabarcoding markers are impractical. As such, monitoring of specific planktonic threats by molecular means may be better achieved by targeted —for example qPCR (Bridle et al., 2010; Krolicka et al., 2022) —and semi-targeted —e.g. clade-specific— metabarcoding approaches (Dario et al., 2017). Target and non-target species ID could also be approached by using multiplexing primers of 18S, cytochrome and other housekeeping markers. However, it is clear from our data that the discovery phase in this area is far from over and metabarcoding still has a role.

In this study, we sampled two aquaculture sites intensively over a single growing season using microscopy and metabarcoding. We aimed to achieve unbiased planktonic sampling and then applied rigorous statistical models to link different planktonic taxa to different salmon health outcomes. Crucially, respective planktonic exposure profiles were both divergent and idiosyncratic, as were the apparent biological drivers of poor gill health. Aquaculture site characteristics, sheltered or exposed, may account for some of the differences observed. However, to make generalisations about the extent of the biological threats to salmonid aquaculture further ‘unbiased’ studies at multiple sites across multiple production cycles are required as well as improving the resolution of the eDNA metabarcoding approach. In parallel, to better understand the mechanisms of gill damage and mortality and which organisms drive them, a more detailed monitoring of salmon gill health is required (e.g. gene expression, histopathology) to help disentangle from the role of direct microalgae toxicity and help set abundance threshold for mitigation purposes. Indeed, new understandings will lead to new approaches for mitigation. These could include protection barriers, aeration technology, early warning signals, new medicinal interventions, and, eventually, methods to enhance fish and gill resilience through functional feeds or selective breeding programs that will enable salmonid aquaculture to thrive in a continuously changing climate.

## ACKNOWLEDGEMENTS

We want to acknowledge our aquaculture partners, who have provided in-kind support to make this study possible; special thanks to staff working on site for their dedicated efforts during sampling period. Thanks also to Dr Owen Wangensteen (University of Barcelona; The Arctic University of Norway) for his support for our data curation using MJOLNIR. We also want to thank Dr Cristian Cañestro (University of Barcelona) for his support in understanding the complexity of *O. dioica* CO1 tracking in our eDNA samples, and Dr Kypher Shreves (University of Aberdeen) for sharing his z-score calculator for the mortality analysis. Acknowledgements also to Dr Margarita Machairopoulou (Marine Scotland) for the nice conversations on zooplankton off West Scotland taxa identification.

Financial support for this study was provided by BBSRC accelerator fund for project “Developing an eDNA approach to monitor planktonic threats to salmonid aquaculture AQUASCAN”, by the Scottish Aquaculture and Innovation Centre (PhD Studentship, Toni Dwyer), and The Royal Society of Edinburgh (Award Ref. Number: 1968).

## Supplementary material

**Figure S1.**
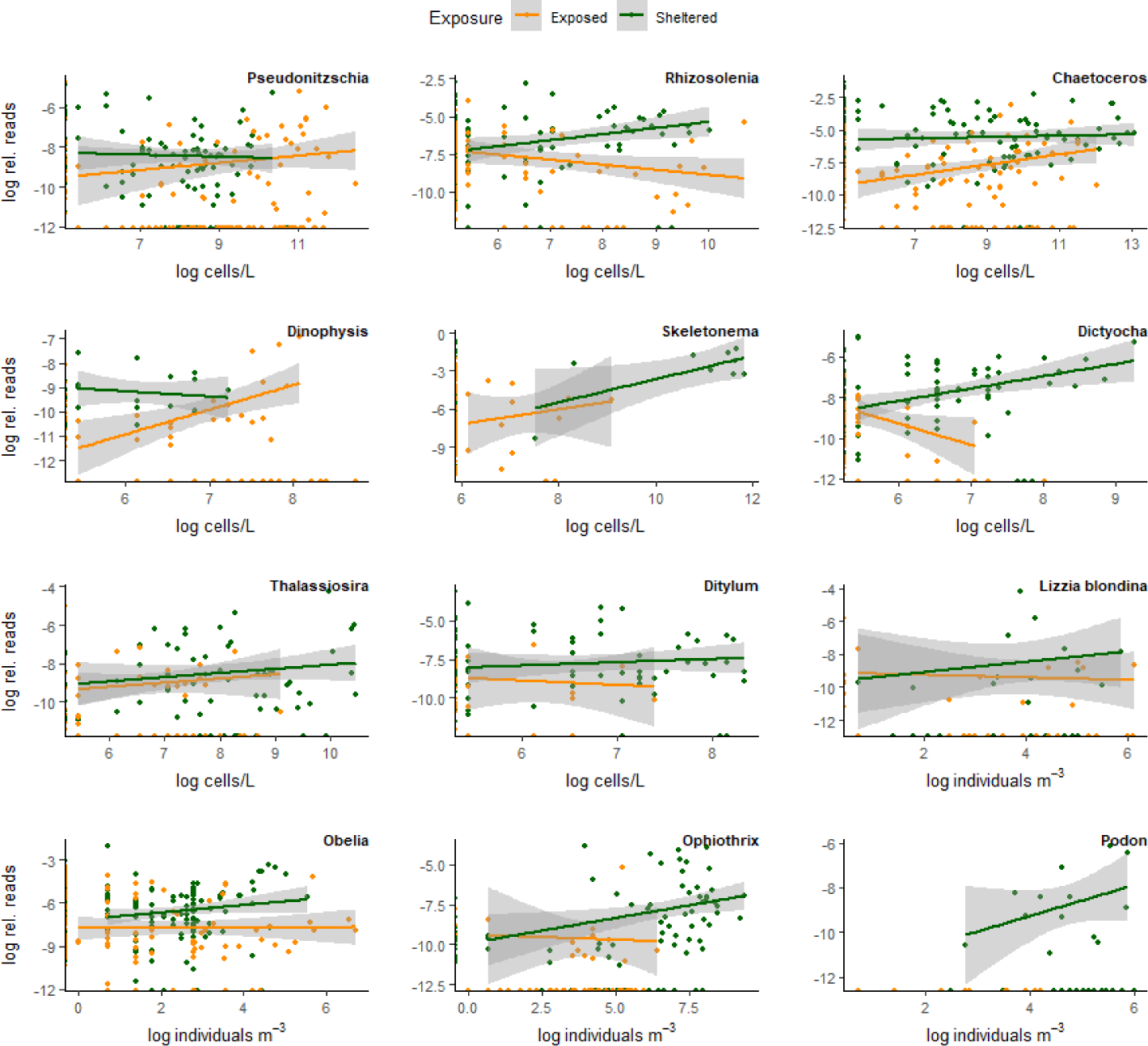
Scatterplots of the relationship between plankton genera identified by eDNA (log of relative reads) and microscopy (log abundance). Analysis is restricted only to genera that were identified by both methods. For each genus, see accompanying Table 2 for regression coefficients and the percentage of discrepancy between the eDNA and microscopy methods (false negatives and positives).

**Figure S2.**
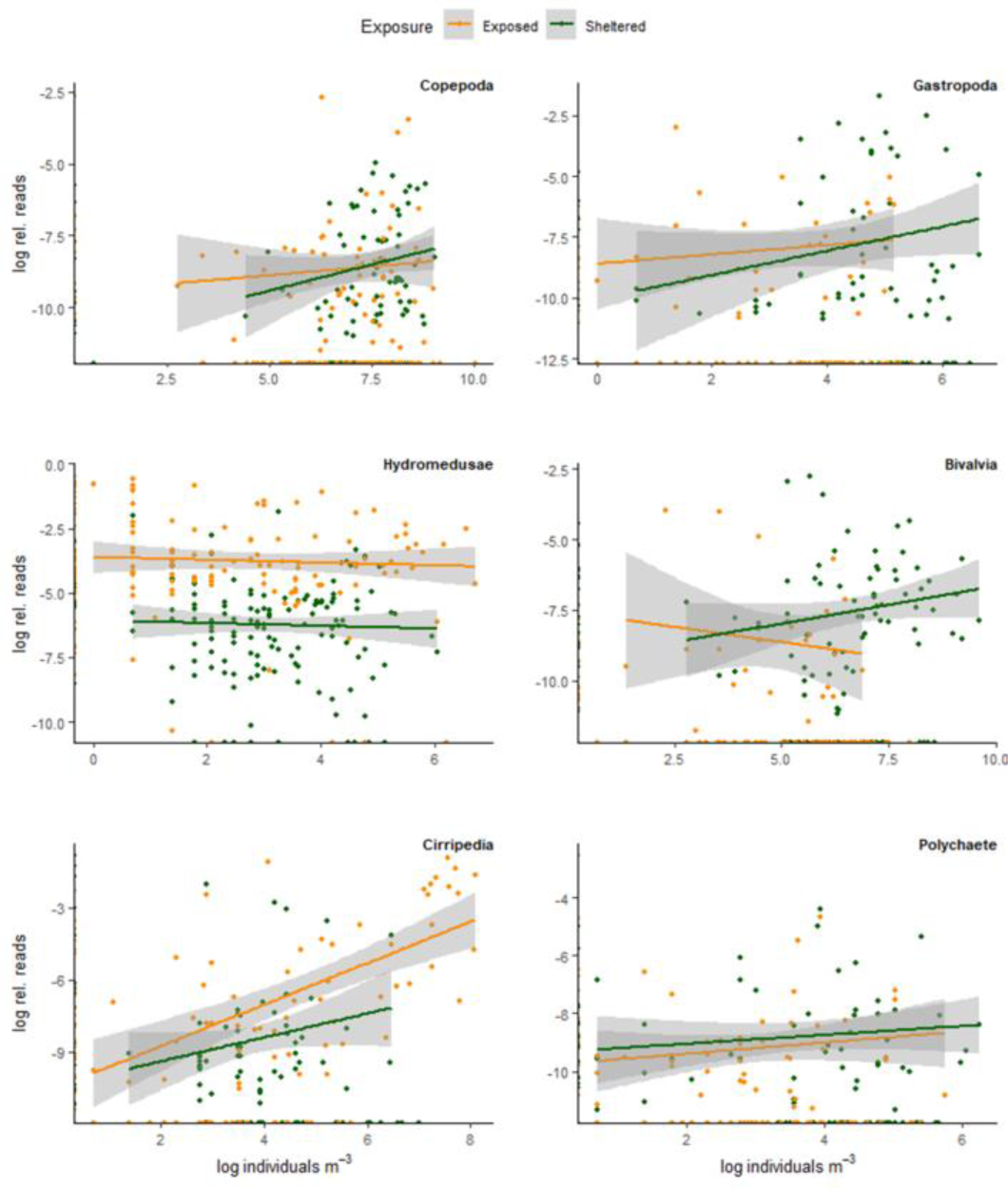
Scatterplots of the relationship between main zooplankton taxa identified by eDNA (log of relative reads) and microscopy (log abundance). Analysis is restricted only to taxa that were identified by both methods (e.g. Appendicularia were not identified by eDNA and are thus excluded). For each taxa, see accompanying Table 2 for regression tests and the percentage of discrepancy between the eDNA and microscopy methods (false negatives and positives).

